# Bacteriophage-Mimetic DNA Origami Needle for Targeted Membrane Penetration and Cytosolic Cargo Delivery

**DOI:** 10.1101/2025.06.16.659898

**Authors:** Anirban Samanta, Mette Galsgaard Malle, Emily Tsang, Marjan Omer, Mads K. Skaanning, Jørgen Kjems, Kurt V. Gothelf

**Affiliations:** Interdisciplinary Nanoscience Center (iNANO), Aarhus University, 8000 Aarhus C, Denmark; Department of Chemistry, Aarhus University, 8000 Aarhus C, Denmark; Department of Molecular Biology and Genetics, Aarhus University, 8000 Aarhus C, Denmark

## Abstract

Inspired by the natural ability of bacteriophages to deliver genetic material directly into host cells, we employed a bottom-up approach to construct a multifunctional synthetic DNA origami needle-like structure. This origami is functionalized with trastuzumab antibodies, cholesterol, protective polymers, and two dyes, which together enable selective targeting and insertion into SKBR3 cancer cells. A disulfide-linked dye payload was attached to the apex of the needle, allowing controlled release in the cytoplasm triggered by the high intracellular glutathione concentration. Real-time tracking of the payload confirmed both successful targeting of the origami structure and subsequent direct cytosolic delivery. By mimicking fundamental mechanisms of bacteriophages, we propose that this artificial needle structure can serve as a prototypical device for the targeted delivery of small-molecule drugs directly into the cytosol.

## Introduction

Bacteriophages, the most abundant biological entities on Earth, have evolved sophisticated mechanisms to deliver their genetic material into host cells^1^. The T4 bacteriophage, for instance, uses its elongated tail fibers to bind to specific bacterial receptors^2^. Subsequently, it deploys a needle-like structure that penetrates the bacterial membrane, a process facilitated by lytic enzymes that degrade the cell wall. This process allows the phage to inject its DNA directly into the cytosol of the host cell. Studying this mechanism can provide inspiration for new delivery devices capable of direct delivery in human cells. This has the potential to solve the major bottleneck in delivery of RNA-based therapeutics which is endosomal escape. It is for instance estimated that only approximately 2% of the RNA payload is delivered to the cytosolusing lipid nanoparticle (LNP) due to endosomal entrapment or exocytosis^3–5^.

To mimic the penetration process of bacteriophage delivery system, we employed DNA origami to construct a needle-like structure (Fig. 1)^6,7^. The DNA origami design was based on a previously reported synthetic DNA-based membrane channels, which used lipidated DNA structures to mimic biological ion channels^8–16^. Although a recent study demonstrated that programmable DNA nanodrills can attach to plasma membranes and induce cell death in a pH-controlled manner^17^, the efficient and controlled insertion of DNA origami pores in the complex membrane of live cells remains a major challenge, which we are addressing here.

**Figure 1.**
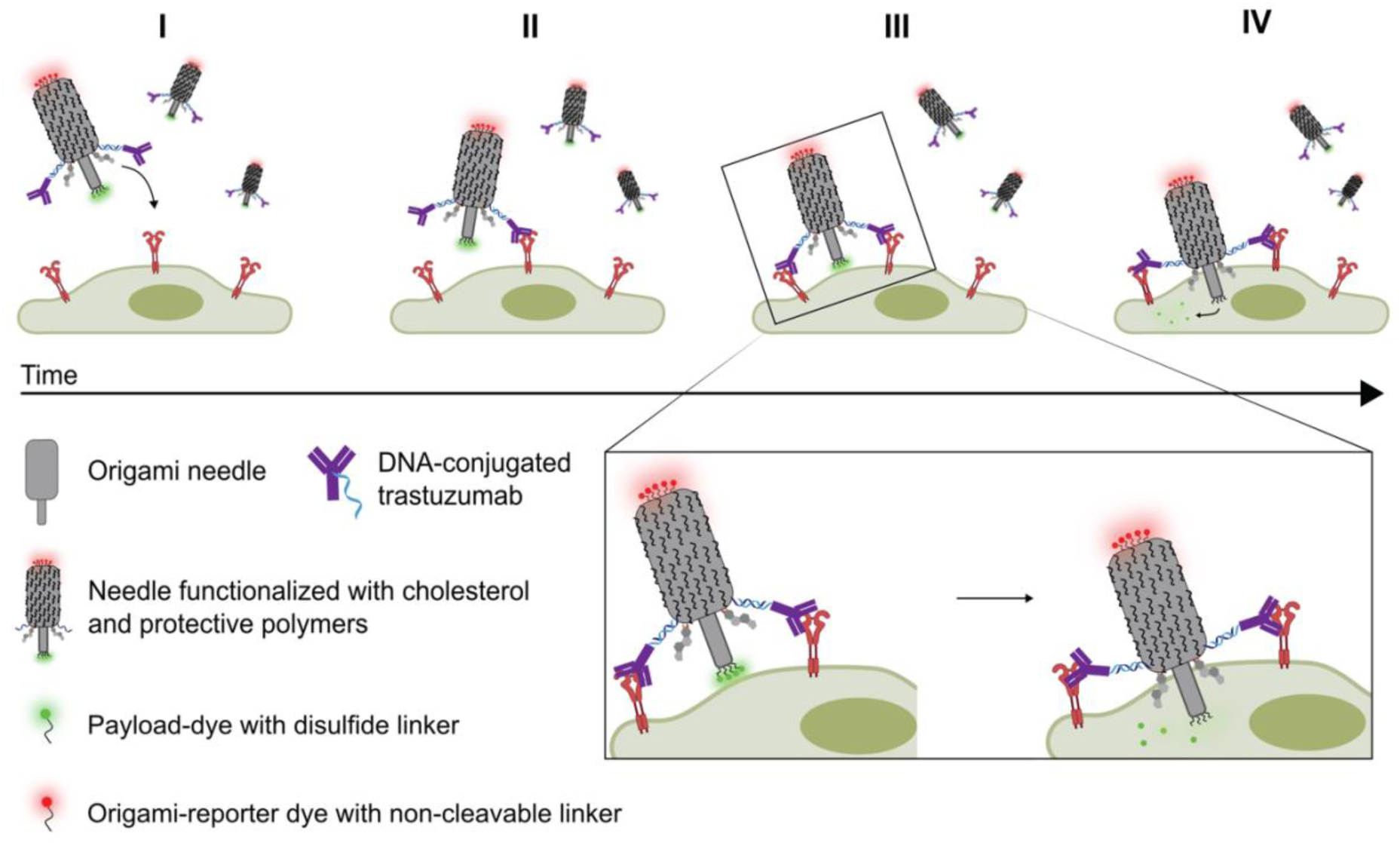
Schematic representation of the DNA origami needle-like structure for cell-specific recognition and direct delivery into the cytosol. The DNA origami needle is functionalized with DNA-conjugated cholesterol and DNA-conjugated trastuzumab antibodies to specifically target human epidermal growth factor receptors 2 (HER2 receptors). The initial specific antibody interaction facilitates membrane association followed by piercing through the cellular membrane with cholesterol mediated insertion (II-III). The cytosolic mM concentration of GSH causes reduction of the disulfide bond, releasing a payload-dye (green) in the cytoplasm (IV), while a red dye attached to staple extensions at the needle base enables simultaneous imaging of the DNA origami.

To facilitate targeted membrane penetration and cytosolic delivery, several components were incorporated into the multifunctional needle-like structure. Firstly, we utilized trastuzumab, an antibody, to specifically targets the human epidermal growth factor receptors 2 (HER2), which are commonly overexpressed in certain breast cancer cells, thereby ensuring selective targeting. Secondly, the structures utilized cholesterol-modified sequences to drive the membrane insertion and penetration process^12,16^. Lastly, the needle-like structures were coated with a biodegradable cationic polymer to protect the DNA from enzymatic degradation in serum^18^.

Upon insertion, the structure releases four fluorescent dyes conjugated to the apex of the inserted needle via disulfide-bonds that are cleaved by the significantly elevated intracellular concentration of glutathione (GSH) in cancer cells at 2-10 mM^19^. The fluorescent cargo was visualized using real-time highly inclined and laminated optical sheet (HILO) microscopy^20^, where single-structures and single-dyes were tracked to determine the intracellular fate. Testing the needle on live cancer cells, we observed selective targeting and cytosolic delivery of the model drug. To the best of our knowledge, this is the first DNA-based nanostructure that selectively binds to the cell membrane and delivers a payload directly into the cytosol.

### Fabrication and characterization of the DNA origami needle

The needle structure is constructed from two main modules: a barrel-shaped base and an elongated needle stem^10^. For the origami design, a hexagonal DNA lattice arrangement was adopted to generate the intended structure in Cadnano^21^. The structure consists of 54 double helices arranged in a needle-like shape, with the stem composed of only six helices and approximately 12 nm in length from its base (Fig. 2a and Supplementary Fig. 1). The staple crossover density was maximized to impart robustness and reduce ion permeability. The predicted dimension of the base structure is approximately 50 nm × 20 nm. After assembly the needle structure was confirmed by agarose gel electrophoresis (AGE) and negatively stained transmission electron microscopy (nsTEM) (Fig. 2b,c and Supplementary Fig. 2).

**Figure 2.**
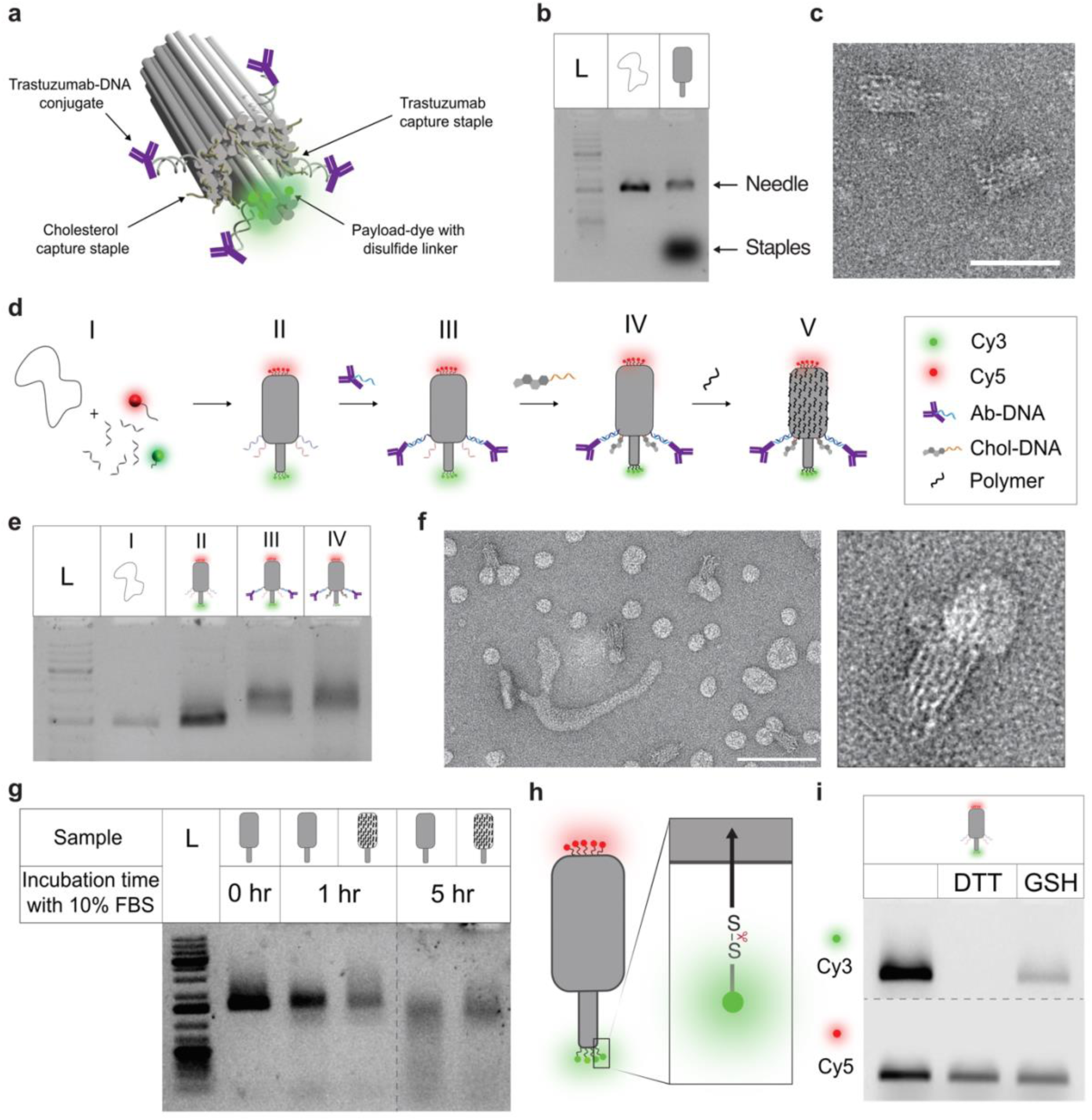
Characterization of a multifunctional DNA origami needle. a) 3D rendering of the designed DNA origami needle consisting of 54 double helices arranged in a hexagonal lattice. b) AGE of the M13mp18 scaffold and the assembled, unpurified origami needle. c) TEM images of assembled origami. Scale bar 50 nm. d) Schematic depicting step-by-step assembly of the needle construct, (I) annealing of the basic needle-like nanostructure from a scaffold and numerous staple strands, including Cy3 and Cy5 dye modified staples and capturing staple strands extensions. (II) functionalization with DNA conjugated Trastuzumab by hybridizing to complementary capturing strands (III) functionalization with cholesterol-modified DNA (IV) adsorption of the cationic polymer onto the negatively charged nanostructure. e). AGE analysis of the needle construct as it is progressively being modified with other entities. (L) 1kb DNA ladder, lane (I) M13mp18 scaffold, (II) purified needle origami (III) Trastuzumab-attached to the origami (IV) Cholesterol- and Trastuzumab-functionalized structure. f) TEM micrograph shows the cholesterol-attached needle origami interacting with POPC derived liposome. Scale bar 100 nm. g) AGE analysis assessing the stability of the needle origami with PCD (N/P = 1) in 10% FBS after 1 hour and 5 hours. h). Simplified schematic of the needle design showing the position of the cleavable disulfide linked payload. i) Gel electrophoretic analysis shows the cleavage of disulfide-linked Cy3 from the origami by reducing agent like GSH and DTT.

To functionalize the needle structure several additions were made (Fig. 2d). First, the needle was functionalized with the antibody trastuzumab-DNA conjugates (Ab-DNA), which bind to four capture strands protruding from the needle (see Fig. 2a and Supplementary Fig. 3)^21^. Functionalization using Ab-DNA on the origami structure was a two-step process. First, a DNA strand was conjugated to the human monoclonal antibody by a lysine directed labeling reaction (LDLR), as previously reported^22^, producing a mono-labeled Ab-DNA conjugate. This is important, as multi-labeling could induce crosslinking of needle structures. The Ab-DNA conjugates were successfully linked to the needle structure in step II-III (Fig. 2d) as confirmed from AGE analysis (Fig. 2e and Supplementary Fig. 4).

Next, the needle structure was functionalized with cholesterol. A total of 17 sites were designed to include staple extensions complementary to cholesterol-bearing strands (Fig. 2d step III-IV; Supplementary Fig. 3). To avoid aggregation, we extended the ssDNA overhangs next to the hydrophobic tag, while maintaining the origami structure’s ability to insert into the lipid membrane^23^. AGE analysis of the cholesterol-appended origami revealed a shift indicating successful attachment of both the Ab-DNA and cholesterol, but with a slight trailing signal, potentially indicating the presence of higher-order structures within the mixture. (Fig. 2e). Initially, we tested the ability of the cholesterol-functionalized origami to insert into synthetic membranes. Here, the needle structures were mixed with small unilamellar vesicle (SUV) and characterized by TEM. The images verify the ability of the origami to insert into the membranes (Fig. 2f and Supplementary Fig. 5).

A major obstacle in applying DNA origami structures in cell media and *in vivo* is their poor biostability at physiological Mg(II) concentrations^24^ and their susceptibility to nucleases in biological fluids^25,26^. Recently, we demonstrated the use of a cationic polymer, poly(cystaminebisacrylamide-1,6-diaminohexane) (PCD), to protect DNA origami under physiological conditions while maintaining structural integrity and addressability^18^. In addition it stabilizes DNA structures by shielding the negative charges. A key feature of PCD is the disulfide bridges in the backbone, which are cleaved in a reducing environment. This lowers the toxicity generally associated with polycationic species^27^. Furthermore, it was demonstrated that PCD, in addition to protection against nucleases, enhanced the association with the cell membrane. To evaluate the optimum PCD concentration, the origami was incubated at varying PCD concentration at room temperature and analyzed using AGE and TEM (Supplementary Fig. 6). TEM analysis of PCD-protected needle structures showed no obvious difference from the unprotected structure at low needle/PCD (N/P) ratio (≤ 1) while aggregation was observed at higher N/P ratios (Supplementary Fig. 6b). Incubating the polymer-protected structure (N/P ratio = 1) at 37 °C in McCoy’s 5A medium containing 10% FBS showed enhanced stability compared to the unprotected structures with minimal sign of degradation even after 5 hours (Fig. 2g and Supplementary Fig. 7). For a more detailed schematic and the molar ratios of each entity added, see Supplementary Fig. 3.

To enable fluorescence visualization, five Cy5-DNA strands were hybridized to staple extensions at the base of the needle (Fig. 2h, red), while four Cy3 dyes were attached at the apex via disulfide bonds using a two-step conjugation process (Supplementary Fig. 8). This cleavable linkage allows release of the Cy3-labeled payload in the cytosol under reducing conditions. Cleavage efficiency was tested using 1 mM dithiothreitol (DTT) and 1 mM glutathione (GSH), which led to a marked decrease in Cy3 signal without affecting Cy5 fluorescence (Fig. 2i), confirming selective disulfide bond reduction and successful release in a complex origami environment.

### A cell-specific and non-toxic payload delivery

Having established a fully functionalized origami needle, we tested the cell targeting, membrane insertion, and subsequent payload delivery in a cell assay using HER2 positive SKBR3 cells (Fig. 3a). Here, we expected a red signal from the Cy5 dyes, positioned at the base of the needle, colocalizing with the cell membrane and whereas the green Cy3 signal was expected to be positioned inside the cells, upon disulfide cleavage in the cytosol. Representative spinning disk confocal microscopy images of the SKBR3 cells, treated with 20 nM of needle constructs and incubated for 2 hours are displayed in Fig. 3b. Sequentially fluorescence monitoring of the Cy3 and Cy5 was conducted with and without overlaying nuclear stain DAPI (blue channel). We systematically investigated the impact of omitting cholesterol, Ab-DNA and PDC resulting in a total of five different configurations and a control sample.

**Figure 3.**
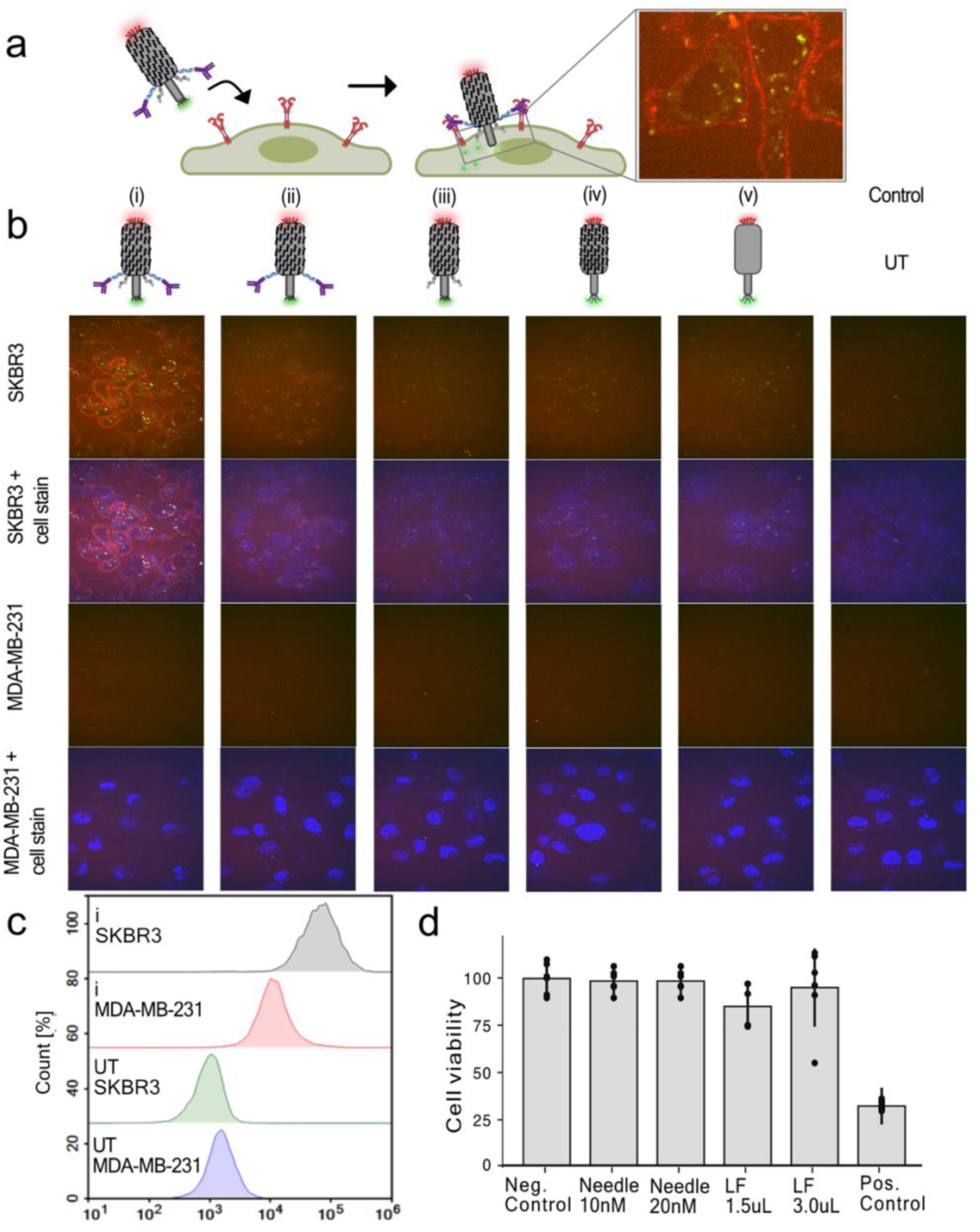
Cell-specific cytosolic payload delivery through a fully functionalized needle construct. a) Schematic representation showing the targeting of the needle construct labeled with a Cy5 needle reporter dye (red) and a cleavable Cy3 payload dye (green). Zoomed-in of micrograph shows the membrane attachment of the red needle dye and cleaved intracellular delivery of the cleavable green payload dye. b) Spinning disk confocal images of SKR3 cells systematically treated with five different needle constructs and control; i) the full construct, ii) the full construct without cholesterol, iii) the full construct without Ab-DNA, iv) the needle construct with PCD, v) the needle construct unprotected and a untreated (UT) control with no construct. Images reflect a significant targeting and payload delivery, completely dependent on functionalizations with both cholesterol and trastuzumab Ab-DNA. Below is shown a control cell line using lower HER2 expressing MDA-MB-231 treated with the five needle constructs showing minimal uptake and targeting. c) Flow cytometry analysis reflects a significant targeted enhancement from the trastuzumab conjugated needle origami with the HER2 positive SKBR3 cells compared to the HER2 negative MDA-MB-231 cells. d) Cell viability of the needle construct after 24 h. No toxicity was found within uncertainties. Lipofectamine (LF) was used as a reference, and positive control shows significant toxicity with treatment of DMSO. The bars represents the mean calculated from n = 6, and the error bars represent ± SD.

From the full construct (i), clear cell targeting is observed, indicated by the red Cy5 signal at the cell membrane. A reduced Cy5 signal is observed for construct (ii) and (iii) omitting cholesterol and Ab-DNA, respectively. The constructs (iv) and (v) exhibited minimal membrane binding, with almost no Cy5 signal, similar to the untreated (UT) control sample. Further controls, using the same series of constructs on a control cell line with low HER2 expression (MDA-MB-231 cells), showed no Cy5 signal from any constructs. Together, this demonstrates the targeting effect of the trastuzumab Ab-DNA together with the necessary membrane recruitment from the cholesterol anchors. Interestingly, the green-channel payload signal was efficiently distributed throughout the cytosol when using the full construct containing both Ab-DNA and cholesterol, indicating enhanced delivery efficiency (construct i, Fig 3B). The majority of the signal in the overlaid images shows only a green signal, although some areas exhibit a yellow signal, indicating colocalization of Cy3 and Cy5. This could suggest uptake of the entire structure, free dyes, or degraded needle structures. However, the green signal appears both as a diffusive weak signal, which could indicate cytosolic delivery, and as large bright aggregates, suggesting endosomal uptake or potential subsequent internal membrane partitioning of Cy3^28^. For additional confocal images, see Supplementary Fig. 9.

Flow cytometry analysis of the needle constructs confirmed specific targeting of HER2 positive cells (SKBR3) with a significant right-shifted signal compared to the control cell line with lower HER2 expression (Fig. 3c). Untreated cells (UT) showed only background signal. Collectively, these methods showcase the initial specific membrane interaction mediated by the trastuzumab Ab-DNA, the crucial membrane stabilization provided by cholesterol moieties, and the added stabilization provided by the cationic polymer, all of which facilitate the cytosolic delivery of the model-payload dyes.

To investigate toxicity from the needle construct, we performed a toxicity assay measuring the cell viability of SKBR3 cells after 24 h, in sextuplicate (*n* = 6, technical replicates) in a 96-well plate, when exposed to 10 nM or 20 nM of the construct (Fig. 3d). As a reference, we tested 1.5 µL and 3 µL lipofectamine 2000 (LF). A positive control was performed using 50% DMSO. We observed no significant toxicity except from the DMSO treated sample.

### Direct observation and tracking of individual payload using HILO microscopy

To investigate the targeting of the needle and uptake of the dyes in HER2 positive cells in more detail, we developed a single-particle tracking (SPT) assay to quantitatively confirm payload delivery of the cleavable dyes into the cellular cytosol. To enable this, the needle structure was labeled with the more stable SeTau-647 dye as the payload conjugated on the cleavable disulfide linker (Fig. 4a). The high photostability, quantum yield, and long lifetime of the payload dye SeTau-647 enable real-time tracking of cytosolic delivery^29,30^. Here, SKRB3 cells were grown on microscopy slides overnight and the membranes were stained before imaging. The media was exchanged to Live Cell Imaging Solution and 9.09 nM origami needle construct was added. Using an Oxford Nanoimager microscope in HILO setting with simultaneous excitation, we performed real-time tracking of both the cell membrane (488 channel) and the payload (640 channel). The HILO setting allows us to track the payload both at the membrane and inside the cells with high signal-to-noise in the focal plane (Fig. 4a)^20^.

**Figure 4.**
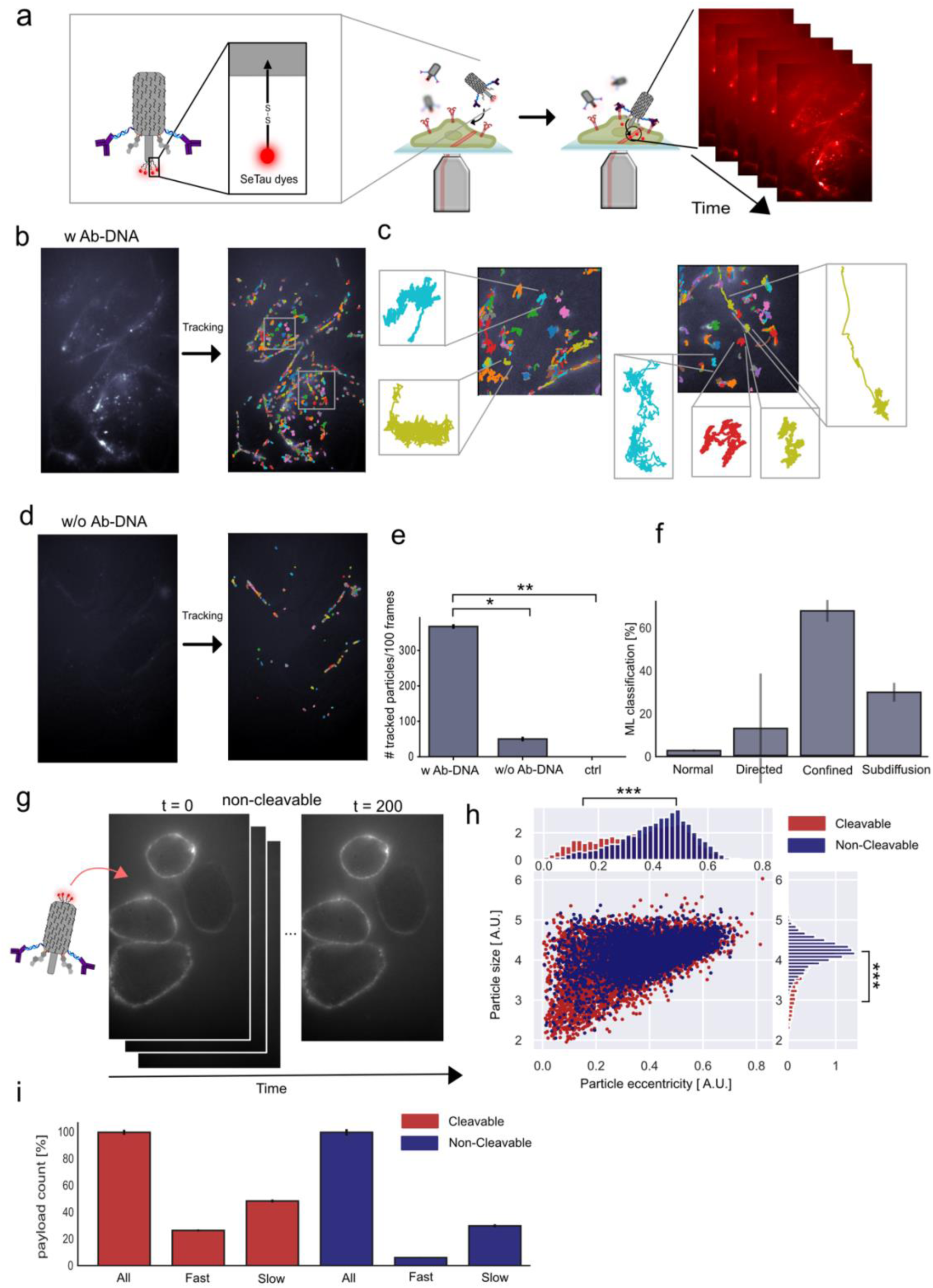
Real-time observation and tracking of individual payload delivery. a) Schematic illustration of the experimental setup. The cleavable disulfide linker on the needle structure was labeled with the stable SeTau-647 dye payload to enable real-time tracking of payload delivery inside cells using HILO microscopy. b) Micrograph showing a representative tracked real-time video of the full needle construct. Colored tracks show the individual identified and tracked payload. c) Zoom-in of individual tracked payloads from the full needle construct inside the cells, depicting different diffusional behavior. d) Micrograph showing a representative tracked real-time video of the needle structure in the absence of Ab-DNA for targeting. e) Quantification of identified and tracked payload using the full needle construct (w Ab-DNA), the construct without antibodies (w/o Ab-DNA), and a control with no construct analyzing the following number oftechnical (n) and biological replicates (N) (n=6, N=5 for w. Ab-DNA, n=3, N=3 for w/o Ab-DNA and ctrl). f) Machine learning classification of tracked payload using DeepSPT algorithm^34^ displaying confined and subdiffusive behavior of the payload (n=9, N=5). g) real-time tracking of needle constructs with non-cleavable seTau-647 staple extensions at the needle base. h) Analyzing the object size and eccentricity of each individual tracked payload from both the cleavable (dyes at the apex) and non-cleavable construct. i) Analyzing the path for each payload from both the cleavable and non-cleavable constructs by tracking both the molecule and cell membrane and classifying payload to be fast (α>0.8) and colocalizing with the cellular membrane and subsequently moving, or slow (α<0.3) and colocalizing with the cellular membrane and subsequently moving. Significance was determined by a two-sided unpaired t-test for e and a two-sided kolmogorov smirnov test for h. Significance levels: not significant (n.s.) indicates P > 0.05, *P≤0.05, **P≤0.01, ***P≤0.001.

To directly investigate the cell-specific cytosolic payload delivery, we first real-time imaged two constructs; one fully functionalized needle construct (w Ab-DNA; Fig. 4b,c) and one lacking the Ab-DNA (w/o Ab-DNA; Fig. 4d). As a control, we tracked cells with no added construct to account for any bleed-through or false-positive signals in the payload channel. All payload was tracked using an in-house developed Python software. The detected payload signal and its corresponding trajectories for the full needle construct are shown in Fig. 4b, where each track is color-coded for clarity in a representative video. We observe a large part of the payload either at the membrane or inside the cytosol. A zoom-in can be seen in Fig. 4c, displaying different diffusional behavior inside the cell from a minor subset of the tracked payload. Importantly, in the absence of Ab-DNA (w/o Ab) (Fig. 4d), significantly fewer payloads are detected, and the signal is predominantly located at the cell membrane, with little to no evidence of cytosolic entry, consistent with the confocal imaging in Fig. 3. For a statistical comparison, we analyzed a minimum of three biological replicates (n=6, N=5 w. Ab-DNA, and n=3, N=3 w/o Ab-DNA and ctrl, with n and N refer to technical and biological replicates, respectively), and, as expected, it revealed significantly higher delivery and membrane targeting in the presence of antibodies and no false-positive for the control with no construct (Fig. 4e). Representative real-time videos are available – see Data availability.

Next, we investigated individual diffusional behavior of the tracked SeTau signal. A spatiotemporal tracking of individual structures interacting with the cell or within the cell reflects the highly heterogeneous behavior and interaction of the molecule^31–33^. Here, we utilized a temporal segmentation model using DeepSPT algorithm^34^ to classify each individual track as either normal diffusive, directed motion, confined motion, or subdiffusive (Fig. 4f). The classification reflects the delivery mechanism, as active transport has previously been described as superdiffusion or directed motion^32,34^, such as endosomal transport. Normal diffusion reflects an unhindered motion, where confined motion characterizes limited spatial interactions confined by boundaries at the cellular membranes. Lastly, a subdiffusive classification might reflect a restrained motion, which could indicate a crowded environment such as the cytosol^33–35^, however it cannot be ruled out that this motion is not endosomal interacting. Analyzing n=9 technical and N=5 biological replicates of tracked SeTau using the DeepSPT machine learning algorithm, revealed that the majority of the tracked SeTau signals were classified as either confined or subdiffusive, consistent with the predominant fraction of the payload being membrane-embedded or cytosolically delivered and moving in crowded enviroments.

To further validate direct cytosolic delivery, we constructed a needle structure with non-cleavable SeTau payloads and performed real-time tracking (Fig. 4g). Here, we observed clear membrane targeting, with only a limited amount of the needle entering the cell. For all detected and tracked signals, in both the cleavable- and non-cleavable needle constructs, we analyzed the size (the radius of gyration of the gaussian-like signal) of the detected particle signal against the measured particle eccentricity (Fig. 4h). Here, the size for a single SeTau payload is expected to smaller than the signal from non-cleaved or membrane-embedded payloads, which consist of four dyes. The eccentricity, which describes the roundness of the tracked individual payload, is expected to be closer to 0, indicating a more circular shape compared to a collection of dyes^36^. As expected, the tracked population of cleaved dyes is both significantly smaller and more eccentric than non-cleavable dyes, with significance levels of p = 1.3·10^-48^ and p = 3.9·10^-64^, respectively (significance tested using a two-sided Kolmogorov Smirnov test). The 2D plot of object size against eccentricity shows that the lower-left population of cleavable dyes does not overlap with the non-cleavable dyes, indicating the delivered payload population. This demonstrates that while a large population of payload are embedded in the membrane, the delivery is selective. Importantly, this suggests that the tracked payload consists of individual delivered molecules rather than entire origami structures.

Lastly, for all tracked payload counts, we fitted the mean-square displacement (MSD) within all trajectories to calculate the anomalous diffusion exponent (α) for each molecule (supplementary Fig. 10)^31,34,37^. Based on this, we classified particles when colocalizing with the cellular membrane and subsequently moving as fast for α>0.8 or as slow for α<0.3. Additionally, particles that remained in the membrane for more than 10 frames with α<0.3 were classified as attached, while particles which were subsequently moving after the 10 frames, were classified as loose. From this, we find the cleavable molecule to have a significantly larger fraction classified as both fast and slow, in accordance with successful cleavage and cytosolic delivery (Fig. 4i).

## Conclusions

We successfully engineered a bacteriophage-inspired DNA origami needle for cell-specific targeting, membrane penetration, and cytosolic cargo delivery. To facilitate specific targeting of HER2 receptors, overexpressed in certain breast cancer cells, we functionalized the needle construct with trastuzumab antibodies, cholesterol anchors, and a cationic polymer for protection from degradation in physiological environments. Across methods, our results consistently demonstrate that the DNA origami needle selectively targets HER2-expressing SKBR3 cells, with payloads mainly delivered directly into the cytosol. This is facilitated by needle apex penetration of the cellular membrane, and the disulfide-linked payload cleavage by high intracellular concentrations of glutathione.

Other large nanostructures have been engineered with lipid-based functionalizations that enable nonspecific cellular targeting^17,38^, and substantial efforts have been devoted to advancing vaccine development and delivery strategies using DNA origami platforms^39,40^. However, membrane-embedded and cell-specific origami structures have largely been unexplored so far. Here, the synergistic effect of antibodies in combination with cholesterol may serve as a cornerstone for design principles aimed at efficiently targeting and attaching DNA origami to the cell membrane, facilitating cargo delivery into the cytosol. In this study, we directly demonstrate payload delivery through glutathione-induced cleavage; however, this principle could be expanded to incorporate other cell specific stimuli in the cytoplasm like toe-hold displacement by RNA or enzymatic by enzymes or by external signals, such as light^41^ or magnetic fields^42^.

We envision that the creation of this artificial, bacteriophage-inspired DNA origami needle could have applications in synthetic biology and serve as a tool for understanding and recreation of simplified cellular communication processes^43^. Additionally, robotic DNA origami structures ^44^ are being developed, including gate transport of molecules^45^ and modular actuator for membrane protein activation^38^, which could further refine our artificial DNA origami needle concept. Most importantly, the needle serves as a prototype for a new class of devices capable of direct cytosolic delivery; offering a potentially universal mechanism for transporting small molecules, peptides, nucleic acids, or protein therapeutics into the intracellular space. By bypassing endosomal entrapment and harnessing stimuli-responsive release mechanisms, such constructs could provide a modular platform for personalized therapies, precision immunomodulation, or intracellular biosensing. As our understanding of nanoscale membrane interactions deepens, and as programmable DNA origami structures grow increasingly sophisticated, this bacteriophage-inspired needle could become a foundational tool in next-generation nanomedicine.

## Materials and Methods

### Design and assembly of DNA origami

The DNA origami structure was inspired by a previous study^10^, with specific modifications made using cadnano^46^ (https://cadnano.org/). Details of the modifications are outlined in the Supplementary Fig S1. The structures were assembled by using a single-stranded DNA scaffold (type: p7249, Tilibit) at concentrations ranging from 3-35 nM. The scaffold was mixed with a 10-fold molar excess of unmodified staple strands (sourced from Integrated DNA Technologies Inc.) and 5-fold molar excess of in-house fluorophore-modified staple strands when required. The sequences of all oligonucleotides used are provided in Supplementary Table 1. The folding buffer contained 20 mM MgCl2, 50 mM NaCl, and 1x TAE. The structures were annealed by using the following method: 90 °C for 2 minute, 65-25 °C ramp (0.5 °C/15 mins), 25-10 °C ramp (0.3 °C/1 min), and 20 °C hold.

### Synthesis of antibody-DNA conjugates

The preparation of trastuzumab-DNA conjugates followed the protocol Märcher et al^47^. Firstly, amino-modified DNA (15 nmol, sourced from Integrated DNA Technologies Inc.) was mixed with 0.1 mg DBCO-NHS ester (BroadPharm), sodium tetraborate (0.1 M, 15 mL pH 8.5) and 15 mL dry DMF. The sample was incubated at room temperature overnight on a shaker before ethanol precipitation, purification (RP-HPLC) and lyophilization. Next, trastuzumab was mixed with a 5-fold molar excess of in-house synthesised lysine directed labelling reagent (LDLR) probe in EEPS (0.5 M, pH 8.5) and NaCl (100 mM). This mixture was left overnight at room temperature before purification using 30 kDa Amicon Ultra-0.5 mL centrifugal filters (Millipore, Germany). After, the trastuzumab-LDLR product was mixed with the DNA-DBCO-NHS ester. A 2-fold excess of DNA was used in 1x PBS. This mixture was left 17 hours to 2 days at room temperature before purification (RP-HPLC) and buffer exchange using 30 or 100 kDa Amicon Ultra-0.5 ml centrifugal filters (Millipore, Germany). The sequences of all oligonucleotides used are provided in Supplementary Table 2 and 3.

### Synthesis of non-disulfide fluorophore-modified DNA conjugates

Amino-modified DNA (15 nmol, sourced from Integrated DNA Technologies Inc.) was mixed with 0.1 mg Cy3-NHS, Cy5-NHS (BroadPharm), or SeTau647-NHS ester (SETA BioMedicals), sodium tetraborate (0.1 M, 15 mL pH 8.5) and 15 mL dry DMF. The sample was incubated at room temperature overnight on a shaker. The solution was diluted to 100 mL with MQ and then mixed with 250 mL 96% EtOH, 14 mL NaOAc (3 M, pH 5.2) and 1 mL glycogen (20 mg/ml). The sample was flash frozen and centrifuged (14,000 rpm) at 4 °C for 45 mins. The pellet was resuspended in MQ and purified by RP-HPLC with Phenomenex Clarity 3u Oligo-RP 50 mm x 4.6 mm column running a gradient of acetonitrile in TEAA buffer (0.1 M, pH 7). The fractions containing the product were collected and lyophilized before being dissolved in MQ.

### Synthesis of disulfide fluorophore-modified DNA conjugates

To synthesize the DNA-disulfide-amino modified strand, amino-modified DNA (15 nmol, 75 μL, sourced from Integrated DNA Technologies Inc.) was mixed with disuccinimidyl suberate (25 mM, 75 μL in dry DMF), MeCN (75 μL), and TEA (1 μL). The mixture was shaken at room temperature for 30 minutes before adding a 10x molar excess of ethylenediamine in MeCN. The mixture was again shaken at room temperature for 30 minutes. After, the DNA was precipitated in an aqueous solution of NaOAc (3 M, 32,1 μL, pH 5.2), EtOH (583 μL), and glycogen (20 mg/mL, 1 μL). The solution was flash frozen in liquid nitrogen and centrifuged (14,000 rpm) at 4 °C for 60 mins. The pellet was resuspended in MQ and purified by RP-HPLC and lyophilized. To form the DNA-disulfide-fluorophore strand, the product 15 nmol in 50 μL) was mixed with Cy3-NHS or SeTau647-NHS ester (234 nmol, 75 μL in dry DMF), MeCN (75 μL) and TEA (1 μL). The mixture was shaken at room temperature overnight before precipitation, purification (RP-HPLC) and lyophilization.

### Assembly of the full construct

To assemble the final construct, the needles (containing the corresponding fluorophores) were annealed and purified from excess staples. The structures were incubated with the antibody-DNA strand for 5-6 hours at room temperature; a 2-fold molar excess per antibody binding site was used. After, the structures were incubated with the cholesterol-DNA strand (sourced from Integrated DNA Technologies Inc.) overnight at room temperature; a 3-fold molar excess per cholesterol binding site was used. Lastly, the in-house synthesized cationic polymer poly(cystaminebisacrylamide-1,6-diaminohexane) – PCD^18^) was incubated with the structures to form a final N/P ratio of 1 (using a 1 mg/ml stock). The structures were incubated for 30-60 minutes at room temperature. All sequences for functionalized oligonucleotides are provided in Supplementary Table 2-10.

### DNA origami purification

The folded needles were purified from the excess staples with 50kDa or 100 kDa Amicon Ultra-0.5 mL centrifugal filters (Millipore, Germany). The filters were filled with buffer (20 mM MgCl2, 50 mM NaCl, 1x TAE or 12.5 mM MgCl2, 50 mM NaCl, 1x TAE or 50 mM HEPES, 0.8 mM MgCl_2_, 0.9 mM CaCl_2_, 200 mM NaCl, pH 7,4 or 50 mM HEPES, 12.5 mM MgCl_2_) and then centrifuged at 13,000 rpm for 5 minutes. After, the filter was filled with the sample (50-400 μL) and centrifuged at 13,000 rpm for 8-10 minutes. The samples were washed 3-4x with buffer.

### Agarose gel electrophoresis

A 0.5 or 1% agarose gel containing 5 mM, 6.25 mM or 12.5 mM MgCl_2_ with 1x TAE or 1x TBE was used to analyze the integrity of the origami structures. The gels were run with 65 - 90 V and pre-stained with 1x SYBR Safe.

### TEM

The purified sample (5 μL, 6 nM) was deposited onto a glow-discharged carbon-coated grid (400 mesh, Ted Pella) for 3 mins. The grid was blotted, dipped on 8 μL of MQ before blotting again. The grid was immediately treated twice with 2% uranyl formate (4 μL) with a 20 second hold before blotting the second treatment. Imaging was conducted with a Tecnai G2 Spirit TEM, operated at 120 kV.

### Liposome TEM experiments

To form a desired concentration of 6.5 mM (5 mg/mL) liposomes, phosphatidylcholine (200 μL in dry chloroform, 20 mg/mL, sourced from Avanti Polar Lipids Inc.) was mixed with 800 μL chloroform and dried in a rotary evaporator for 20 minutes at 40 °C. After, the sample was put under a high vacuum line for 1 hour. The dried phosphatidylcholine was dissolved in 2 mL of 1x TAE buffer with 10 mM MgCl_2_. The sample was vortexed three times for 10 seconds each before sonicating for 20 minutes at room temperature. The liposomes were mixed with the cholesterol-functionalized needles in a 1/100 dilution for 1 hour at room temperature before TEM analysis.

### Flow cytometry studies

Each well of a 96-well plate was seeded with 50,000 suspended cells (either SKBR3 or MDA-MB-231) and incubated O/N. The following day, the cells were washed in 1x PBS. The needle structrue was mixed with cell media and 20 nM, 200 μL of contruct was added and incuabted at 37°C for 4 hours. For untreated cells, 200 μL of media was added. After incubation the media and needle contruct was removed, and the cells was washed 2x times with PBS. 120 μl of trypsin-EDTA was added, incubated for 8-10 mins, and then 130 μl of media was added. The samples were then transferred to an Eppendorf tube and centrifuged at 2,000 g for 10 min.

### Cytotoxicity studies

Each well of a 96-well plate was seeded with 10,000 suspended SKBR3 cells and incubated O/N. The cells were treated with the needle construct in 2 concentrations (10 nM and 20 nM) and lipofectamine (1.5 μL and 3.0 μL), all in 600 μL total volume, all in sextuplicates (n = 6 technical replicates). After 24 hours of incubation, the cells were washed with 1x PBS and incubated in 10% alamarBlue reagent (Molecular Probes, Life Technologies) in cell media at 37 °C for 2 h. Cell viability was determined by fluorescent intensity measured using a plate reader (FLUOstar OPTIMA, Moritex BioScience) with an excitation wavelength of 540 nm and emission wavelength of 590 nm.

### Acquisition of needle uptake using Spinning Disk Confocal imaging

All confocal imaging were accomplished using a IX83 inverted spinning disk confocal microscope, Olympus, with a 60x water objective with cellVivo CO2 and temperature control at 37°C. Four line laser were used for recording iChroms MLE 405nm, 488nm, 561nm, 640nm with Quad band filters: 440-40 / 521-21 / 607-34 / 700-45. SKBR3 and MDA-MB-231 were seeded in 6 channel micro slides (μ-Slide VI 0.4 ibidi-treat; 80606, ibidi) at a density of 1·105 cells per well in 100 µL of culture medium and incubated overnight. nucleus were stained using Hoechst 33342 (Molecular Probes) diluted 10,000 times and incubated for 10 minutes. 20 nM construct mixed in Live Cell Imaging Solution (InvitrogenTM, A14291DJ) were added and incubated for 4 hours.

### Acquisition of real-time imaging data

All real-time tracking experiments of the origami needle construct were accomplished using an Oxford Nanoimager (Oxford, UK) microscope. The data was acquired with a 100x 1.4NA oil immersion objective at 37°C and the samples were administered with an external peristaltic pumping system with a flow rate of 0.1ml/min. The image dimension for each channel is 428 times 684 pixels with a dynamic range of 16-bit grayscale, recording 2 channels simultaneously. The field of view corresponds to a physical field of view dimensions of 50 µm times 80 µm per channel. All tracking experiments is performed in HILO setting.

### Recording of real-time tracking experiments of the origami needle construct

SKBR3 cells were seeded in a 6 channel micro slides (μ-Slide VI 0.4 ibidi-treat; ibidi) at a density of 1·105 cells per well in 100 µL of culture medium and incubated overnight before the experiments. After overnight incubation, the cell membrane and nucleus were stained at 37°C for 15 min in culture medium supplemented with Wheat Germ Agglutinin, Alexa Fluor™ 488 Conjugate (Invitrogen, W11261) The origami needle construct was conjugated both with and without trastuzumab antibodies and the disulfide linked payload was SeTau-647 dyes (SETA BioMedicals) for all live tracking experiments. All cells are imaged in bright field. Cells and constructs were further tracked using simultaneous laser excitation with 488nm laser (7%) and 640nm laser (7%) and 200 ms exposure time, where the signal is divided in two channels using the dichroic mirrors on the ONI setup with one channel tracking the disulfide linked SeTau-647 payload (640) and the other channel tracking the cells and the origami reporter dye. The tracking experiments was performed in Live Cell Imaging Solution (InvitrogenTM, A14291DJ) and with 9.09 nM origami construct added.

## Data availability

Supporting videos are available through ERDA at: https://anon.erda.au.dk/sharelink/b59i0cjr1w

All data used for this study will be available through ERDA server upon request.

## Acknowledgements

This work was funded by Novo Nordic Foundation Challenge Center for Multifunctional Biomolecular Drug Design (Grant No. NNF17OC0028070). We are also grateful for the LC-MS spectrometer funded by the Carlsberg Foundation (Grant No. CF21-0579).

M.G.M was funded by Lundbeck Foundation (grants no. R380-2021-1393).

AS is currently at Ramakrishna Mission Vidyamandira, Belur Math, India-711202.

## Author contribution

AS designed the needle and the modified versions of the needle. AS and ET prepared and characterized the full construct and studied interactions with cells in flow cytometry and confocal spectroscopy. MGM designed, conducted and analyzed the tracking experiments and confocal microscopy. MO and MKS optimized the cytosolic delivery. KVG conceived and supervised the project, JK co-supervised the project. AS, KVG, ET and MGM co-wrote the MS and all authors proofread the MS.

^+^These authors contributed equally: Anirban Samanta(AS), Mette Galsgaard Malle (MGM)

## Supporting information

Is available for this paper.

