## Supplementary Information for "Bacteriophage-Mimetic DNA Origami Needle for Targeted Membrane Penetration and Cytosolic Cargo Delivery"

*Affiliation:*

**Figure S1. Blueprint of scaffold and staple routing.**

Illustration of the scaffold and staple routing, as well as the helical arrangement, exported from Cadnano. Helix 0 to Helix 5 makes up the needle tip.

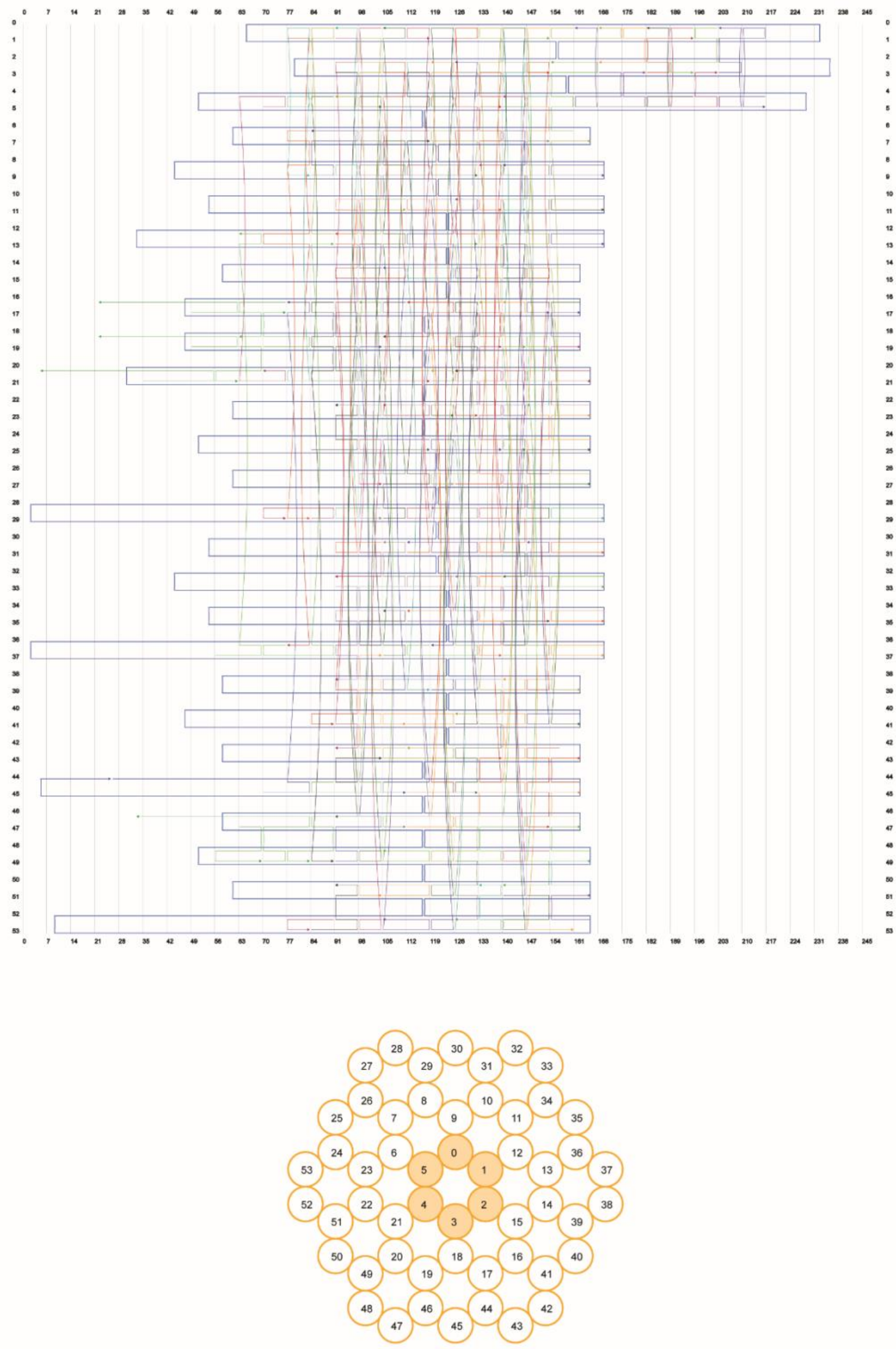

**Figure S2. TEM image of unfunctionalized needle.**

Full TEM image of the purified structure in 20 mM MgCl2, 50 mM NaCl, 1x TAE, 52,000x magnification. Scale bar is 100 nm. A crop-out of this image was used in Fig. 2c.

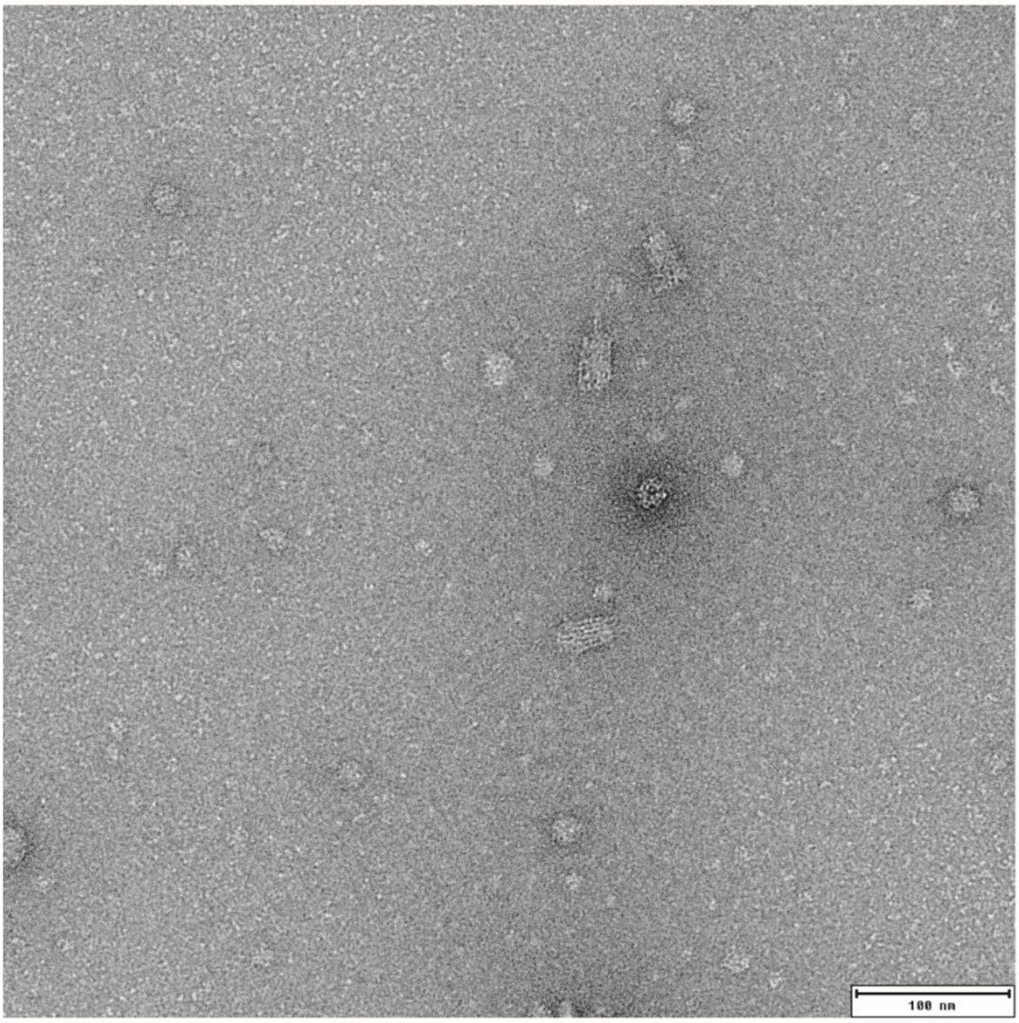

**Figure S3. 3D and simplified illustration of the origami needle design.**

(a) A 3-dimensional rendering of the origami needle design depicturing the estimated dimension of the base structure as 50 nm × 20 nm. b) Schematic of the needle tip of the origami design showing the positions for capturing staple strands extensions for functionalization with both Trastuzumab antibodies and cholesterol. (c) A simplified illustration highlighting the capturing staple for trastuzumab and the cholesterol capture staples, as well as the payload and reporter dye binding configurations. (d) A schematic of the assembly progress of the full needle.

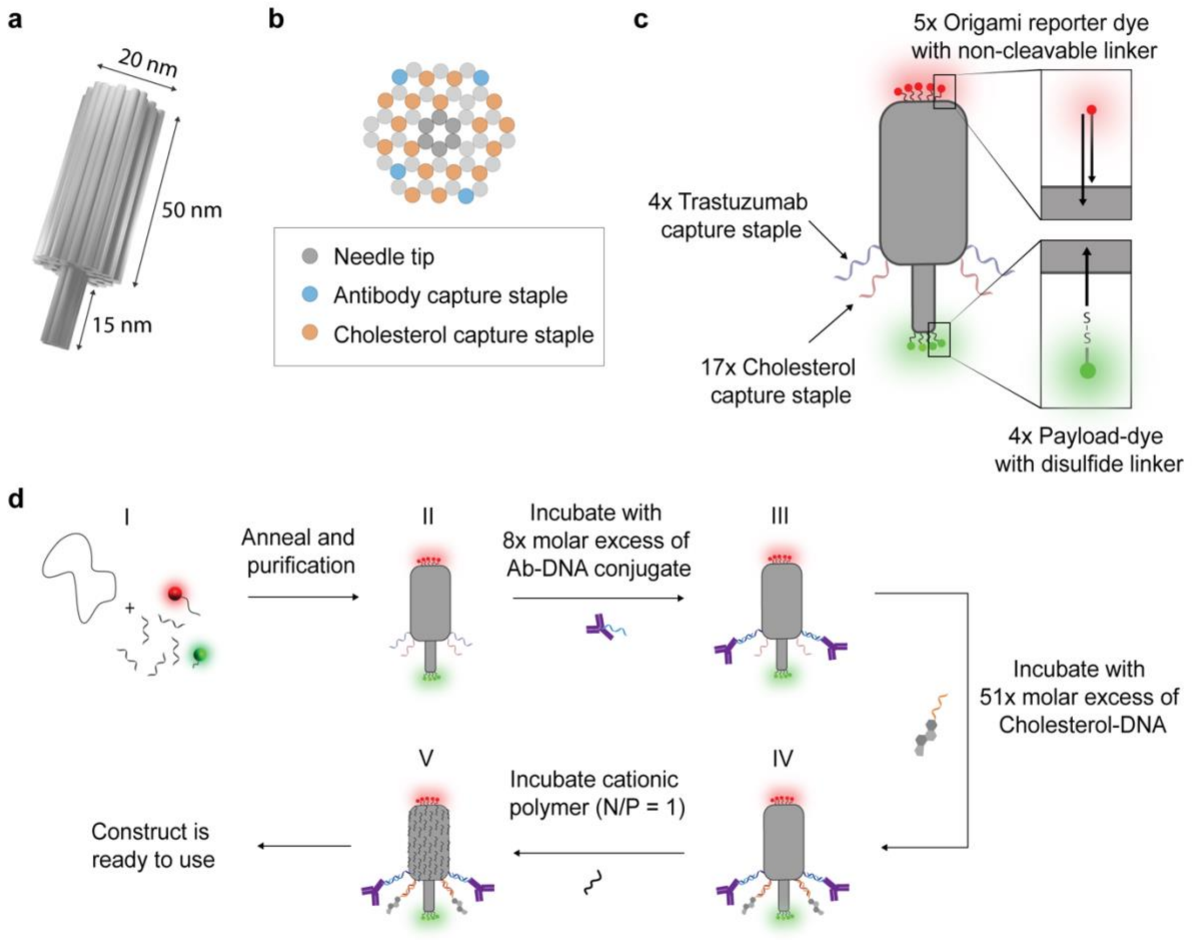

**Figure S4. Functionalization of the needle with trastuzumab-DNA conjugates.**

The origami needles were mixed with varying equivalence of trastuzumab-DNA conjugates at room temperature. After, the samples were analyzed on an agarose gel. The gel conditions were 1% agarose with 12.5 mM MgCl2, 1x TAE, 80V, 45 mins, SYBR safe stain.

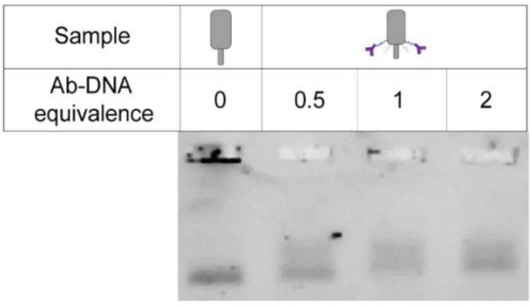

**Figure S5. TEM image of cholesterol-functionalized needles with liposomes.**

The liposomes were mixed with cholesterol-functionalized needles in a 1/100 dilution for 1 hour at room temperature. Magnification is 52,000x and the scale bar is 100 nm. A crop-out of this image was used in Fig. 2f.

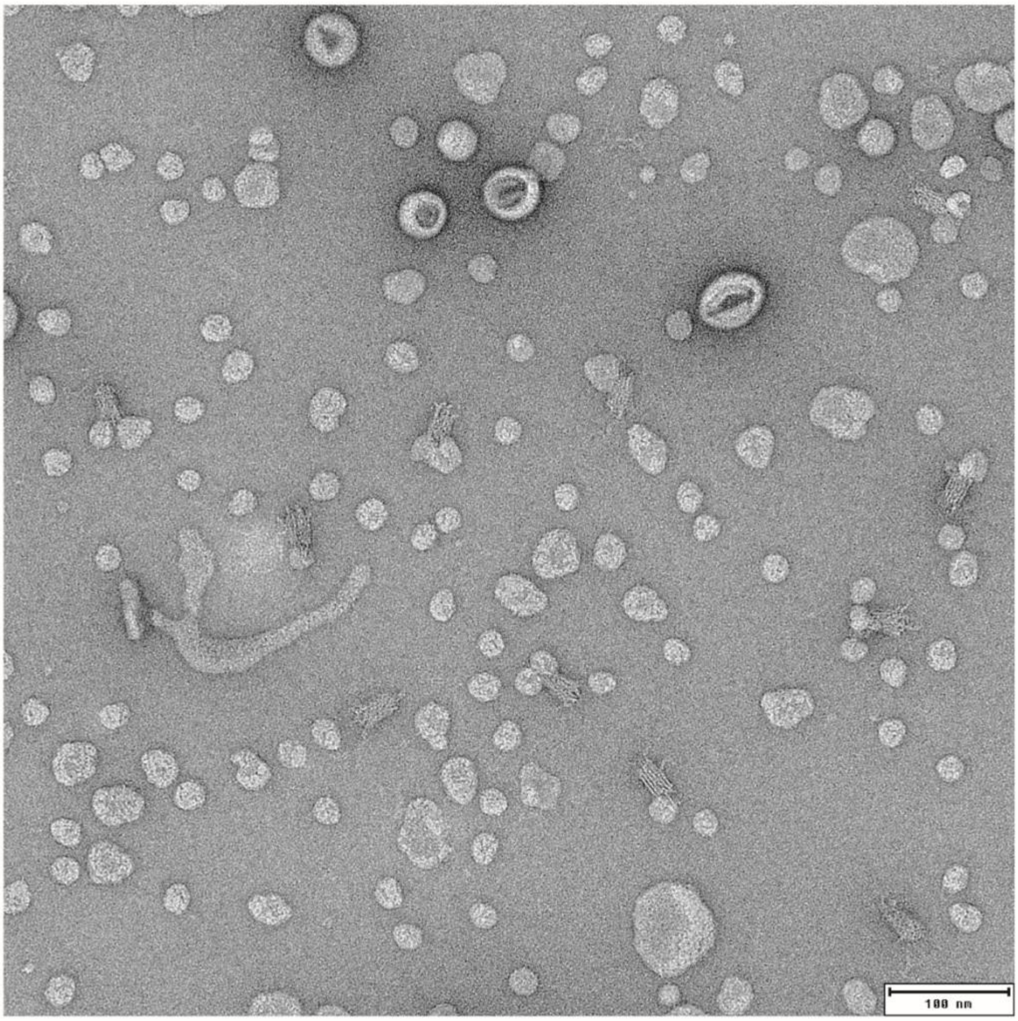

**Figure S6. Optimum PCD concentration for needle contruct protection.**

(a) Agrose gel electrophoresis of the needle origami mixed with different N/P ratios of PCD. The origami was purified into a lower salt buffer (50 mM HEPES, 0.8 mM $MgCl_2$, 0.9 mM $CaCl_2$, 200 mM NaCl, pH 7,4) before mixing with PCD. The gel conditions were 1% agarose with 12.5 mM $MgCl_2$, 1x TAE, 90V, 35 minutes with SYBR safe stain. (b) TEM images of needle structures with the cationic polymer with and N/P ratio of 0, 0.5, 1 and 5. Magnification is 42,000x with a 200 nm scale bar.

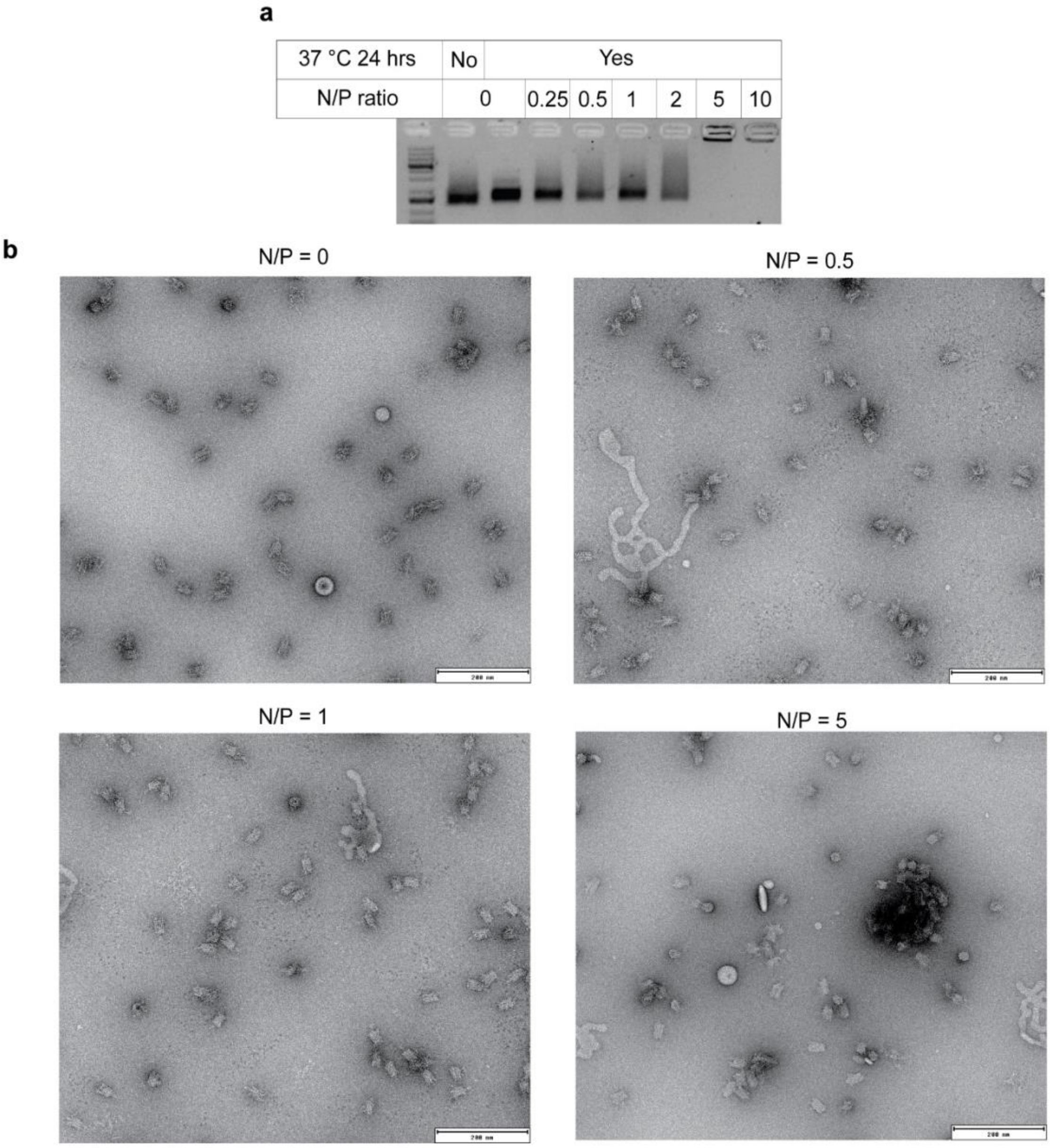

**Figure S7. Stability study of needle protected with PCD polymers in 10% FBS.**

(a) Agrose gel electrophoresis assessing the stability of the needle origami with PCD (N/P = 1) in 10% FBS. The origami was purified into a lower salt buffer (50 mM HEPES, 0.8 mM $MgCl_2$, 0.9 mM $CaCl_2$, 200 mM NaCl, pH 7,4) before mixing with 10% FBS at 37 °C. The gel conditions were 1% agarose with 6.25 mM $MgCl_2$, 1x TAE, 80V, 35 minutes with SYBR safe stain. A crop-out of this gel was used in Fig 2g.

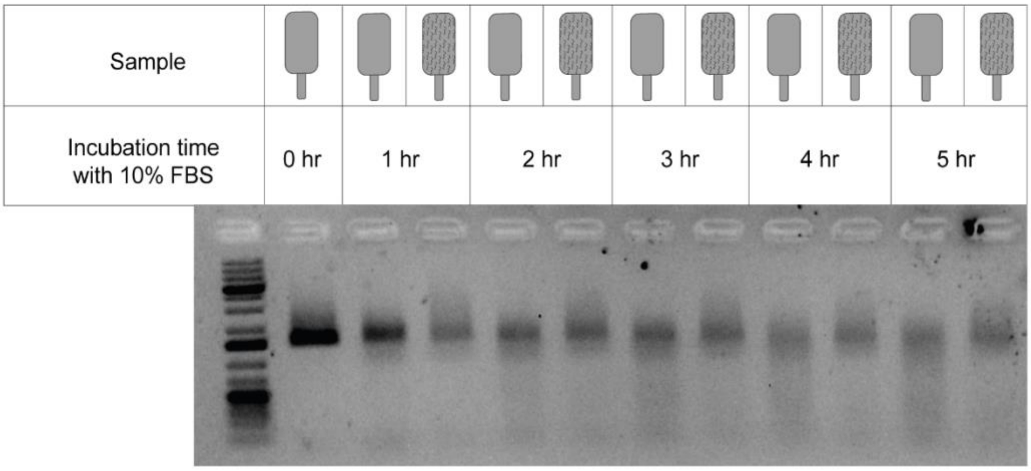


**Figure S8. Reaction scheme for synthesizing the payload-dye with the disulfide cleavable linker**

Amino-modified DNA staple strands was reacted with disuccinimidyl suberate and shaken at room temperature for 30 minutes. The produce was precipitated and HPLC purified before reacting with Cy3-NHS or SeTau647-NHS ester. The mixture was shaken at room temperature overnight before precipitation, purification (RP-HPLC) and lyophilization.

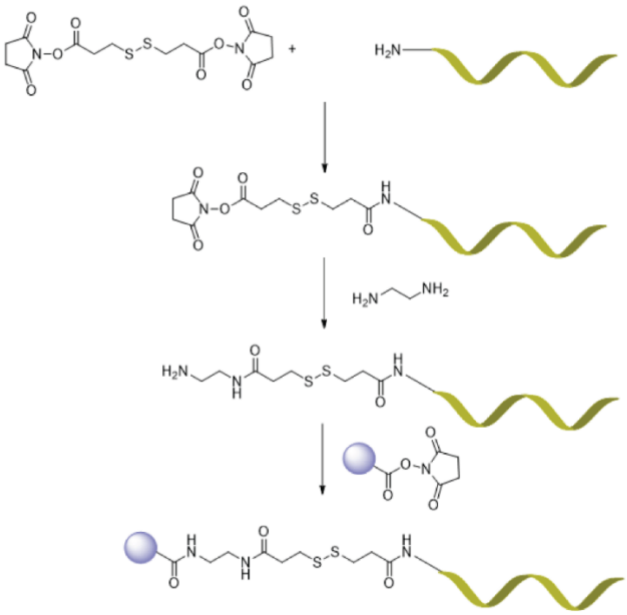

**Figure S9. Needle uptake studie using Spinning disk confocal imaging.** Confocal imaging for uptake studies for five needle constructs and a ctrl with no added construct. The five construct are a) the full construct, b) the full construct without cholesterol, c) the full construct without antibodies, d) the needle construct with PCD, e) the needle construct unprotected. Which allow us to study the effect of antibodies, cholesterol and PCD protection by systematically leaving one out. From the quantitative images it is found that the full construct shows significant more uptake than the construct with no antibodies or cholesterol, indicating a clear targeting effect of having both present and showcasing that the needle structure indeed is capable for a targeted delivery of payload.

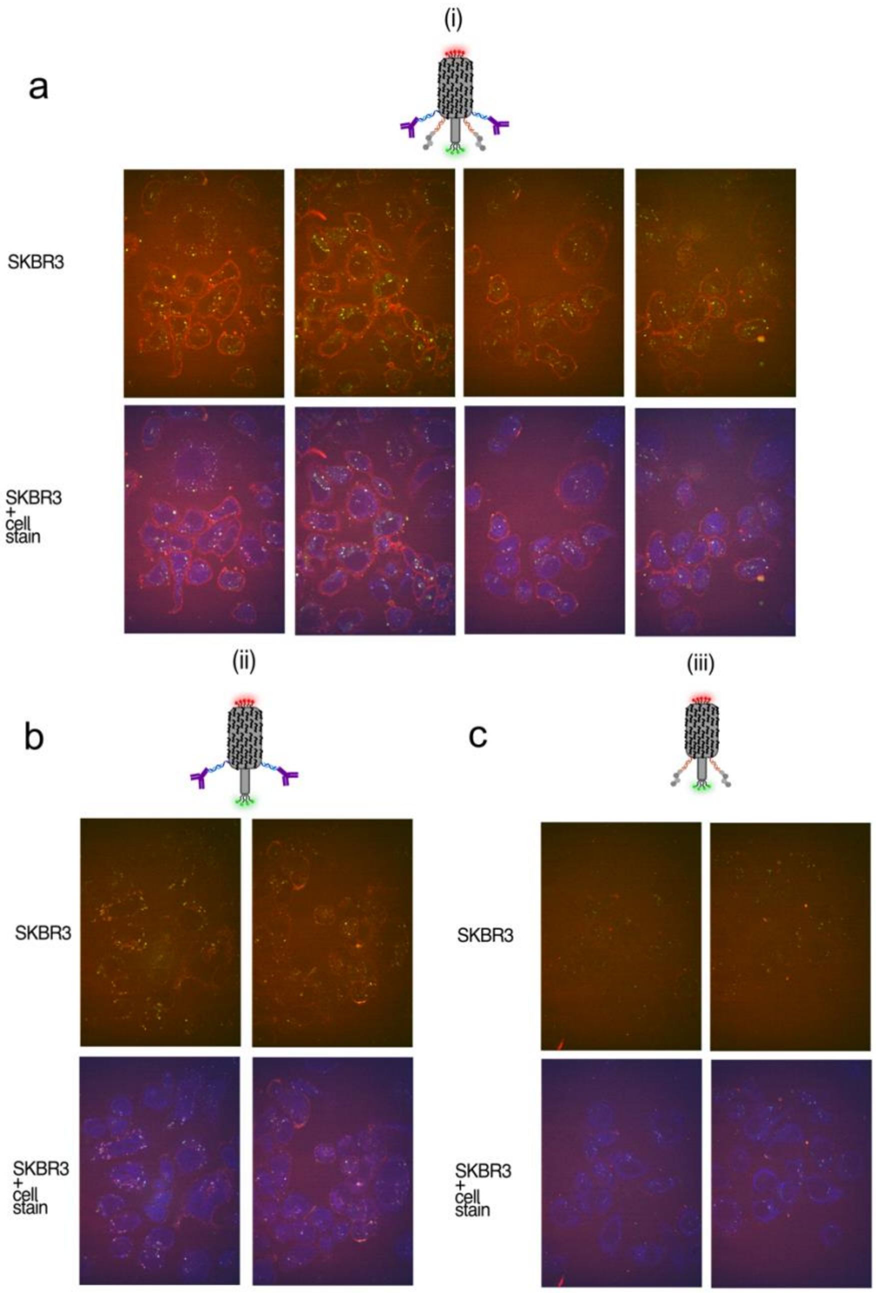

a
(i)
SKBR3
SKBR3 + cell stain
b
(ii)
SKBR3
SKBR3 + cell stain
c
(iii)
SKBR3
SKBR3 + cell stain

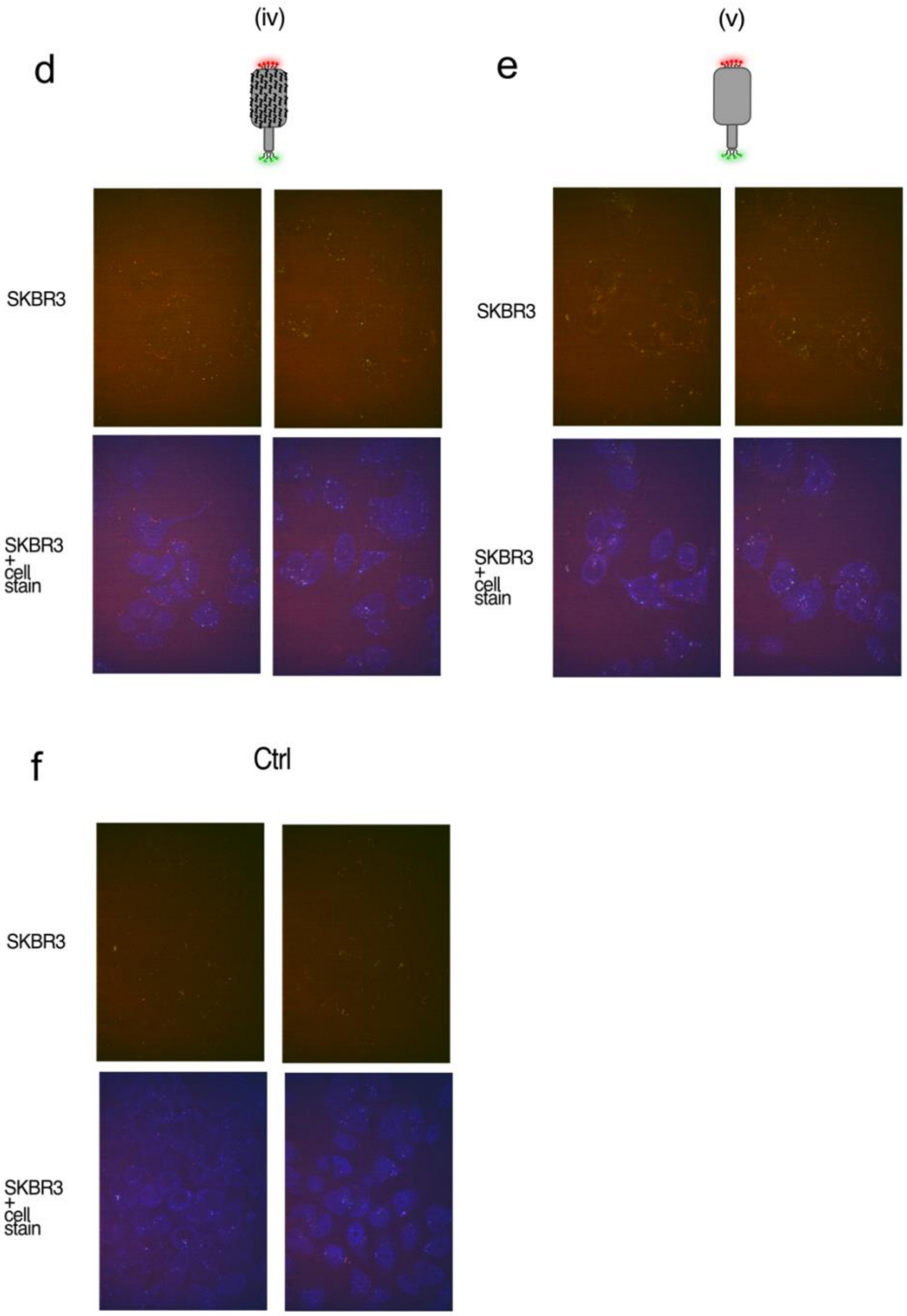

d
(iv)
SKBR3
SKBR3 + cell stain
e
(v)
SKBR3
SKBR3 + cell stain
f
Ctrl
SKBR3
SKBR3 + cell stain

**Figure S10. Single molecule tracking insights of cleavable and non-cleavable linked seTau.**

a) From SPT of needle structure with both cleavable and non-cleavable linked seTau payload we fitted the Mean-square displacement (MSD) within all trajectories to calculate the anomalous diffusion exponent ($\alpha$) for each molecule. The $\alpha$ values are found to be highly significant larger for payload with cleavable linkers than non-cleavable linked payload, which might be due to delivery and not only membrane association. (significance tested with a two-sided Kolmogorov Smirnov test, $P = 4.87 \cdot 10^{-10}$). b) Mean-square displacement (MSD) and anomalous diffusion exponent ($\alpha$) this we determine for each particle if it is fast ($\alpha$>0.8), colocalizing with the cellular membrane and subsequently moving, or slow ($\alpha$<0.3) colocalizing with the cellular membrane and subsequently moving. Moreover, molecules fount in membrane more than 10 frames and $\alpha$<0.3 were classified as attached, and molecules in membrane more than 10 frames, $\alpha$<0.3 and subsequently moving were classified as loose.

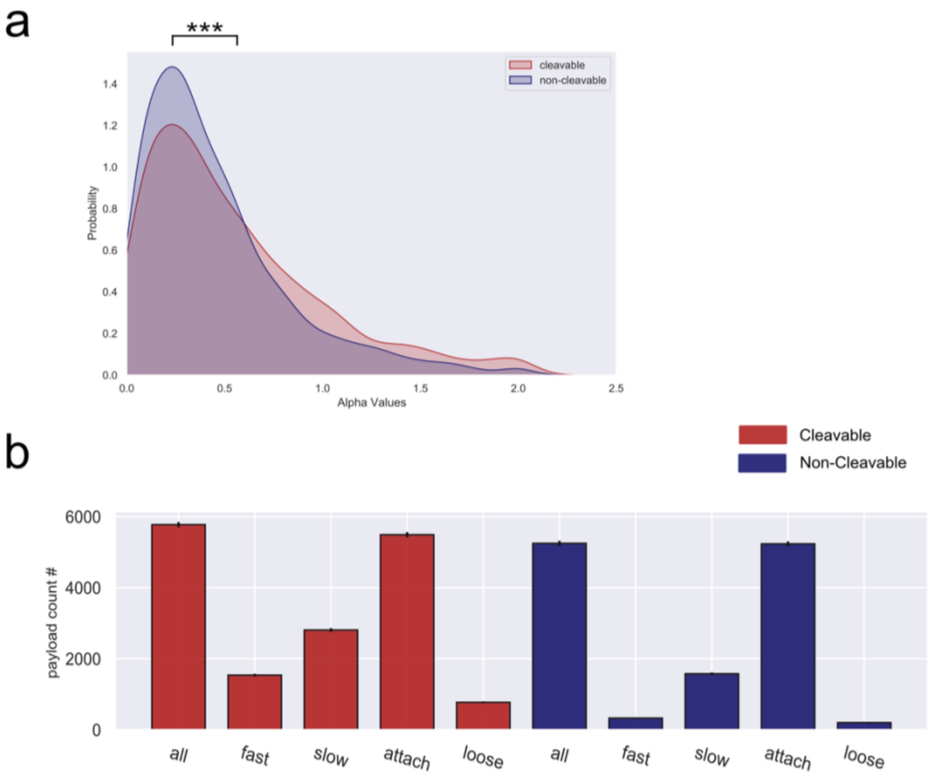

**Table 1. Table of core sequences for constructing DNA origami needle**

| Name | Sequence |
|---|---|
| 0[104] | TCACCGTTTGACCCAACGGAGTATAAGCTAAAATTCGCATTA |
| 0[153] | ATAAGTGGTTTGCCGTTTGCCTTACCAAGAGGTTTTTATTAATTTTAAA |
| 0[160] | CAGGCGGGTAACGGGGTCAGTGCCTTGAGTTTTGC |
| 0[181] | TAGCGGGGTAACAGACAGGAGTGTACTG |
| 0[202] | CCTCAAGGTTAATGTCATACATGGCTTTCTTGATA |
| 1[77] | AACCGCCGGCTGACCTTCATCGCTTGCCCTGACGAGCCATTC |
| 1[91] | GAGCCACGCGTAACACGAACTTACCACATCACCAGAGCAAGC |
| 1[119] | CAAGCCCCAAGAGGACAGATGCTTGAGATGGTTGCAAGCACC |
| 2[118] | CACAGACATTACAGGTAGAGTACTAATTTCAACTTAACTCAC |
| 3[91] | AGACGTTGAGGGGGTAAAAACTCCAGTTCCCTTATAAATCAA |
| 3[126] | GCTAAACTGCGAATCCAAATAAAACAGCATATTCCCACCAGA |
| 3[168] | GAGGCAGCCGCCGCCAGCATTCATAATCTCAGTAC |
| 3[182] | GATTGGCTGATGATTGCCCGTATAAACA |
| 4[90] | GGCTCCAACAGGTCAGGATTACTGACTATTCAGAACAGAACA |
| 4[139] | AAGGAATAACTTTCCCCACAAAACAATGCTTTTTTGTGAATA |
| 4[153] | GAAAGGAGTTTAACCTAATTT |
| 5[70] | TTAAACACCTTTAATTGTATCCCAGACCGGAAGCAAAAGCGG |
| 5[105] | GACAACAACCATGGTTTTCACCTGTATGCCTGCAAAAGAAGT |
| 5[140] | TATTAGCCCGTCGATCACCGTAGGGAAGTCTGTCCGCCAGTA |
| 5[168] | AAAATCACCGGAACACCAGAGGTCAGAC |
| 6[83] | GGCTTGCACCCTCAGAAGTTTGGTAGCT |
| 6[118] | TCCGCCCACGCATATCTGGAAGTTTCTTTTCCTTTTGATAAG |
| 6[125] | GTTTTCAGGAAACGTCACCAAATCAGATATAGACTGAAGTAC |
| 6[160] | GAATCAAATCTTTTGAGAATAAGTGACAGGAGGTTGTAATAAGTTTTTT |
| 8[132] | TAGCAAGGCCGGCTGCACAAGAAAAATAGGTATTAAACCACC |
| 9[84] | TAAAACATCATGAGGCAGCGAAAGACAGTATTCGGTCGCTGA |
| 9[133] | TGAATTAGAGGGTTACCCATGACTACAAAGGAAACCAGATAG |
| 10[111] | AAAGTACCCAGCGACTTTGAGACGAGGGGTAGTAGAAAGGTGGCATCAA |
| 10[125] | AAATTGCTACGACAAAAGGTAGATAAGTCCTGACTGAGAGTC |
| 12[90] | ATAGGCTACCCTCAAGGTTTAACCGATAGTTGCGCAAAAAAA |
| 12[132] | ATTTTGTAATAGGAGATATAAGGTCATAGTAGCGCGCTACAA |
| 12[139] | TAAGTTTTAGCAAATAAGACTTAGGTTGGAGACTACCTTTTT |
| 13[91] | CAGAACGTACCTTAGGGGTGCCAGTCGGGAAACCTTTTCTTT |
| 14[104] | TGTGAATAGTAGTAAAACCAGAAGATCGCACTCCACGGCGGA |
| 15[98] | AACGGAACAACATTAGCCCTCGAATTTTGTTGAAACCTTTAATTGCTTA |
| 15[140] | CGAGGAAGCATGATCGTAGAAGTGATAACATAATTACTAGAA |
| 16[90] | TTCAACTACGACGATAATAGTTAAACAGTTATAGTTATTTTT |
| 16[111] | TTGAGATGCGAGAGGCTTTAAATAGATAGGGTTGAAACTCAA |
| 18[104] | TTTGCCAAGTAAATATAGTTACACCCTCTACAGACAGGGAACCGAACTG |
| 18[146] | GAGATAAAACAGTTAGTACAATACCGTACACGGAACATATGG |
| 18[153] | TATCAGAAAGTCAGTACAGAGAAATAAAGGATTATACTTCTG |
| 19[105] | ATGACCATAAATGAGCCGGCCTTGCTGGATTTACA |
| 20[83] | CAGAAGCAACTCCAAAAGGAGGCTTGATGTACCGCAATACAC |
| 20[118] | TACAAAAATCAGGTCTTTAATGCGCGCCCAAGATTCACCAGT |
| 21[119] | TCCCAATAATAATTCATTTTCGTATAGCAAATTCATTAAAGG |

22[104] AGGTCATTACGGTGACCGATACATCGGAGACTAAATACAAAG
24[118] TTATTCCATATAACACAAACAATTCGACGAGAGCCAGCAGCA
24[132] AAGATTAATTAAATCCTTTGCAAGAAACTGATTATAATAATG
24[139] TTAAATCTTTTATCACAAAATAGAAACGCATAAAAGAACACC
26[146] CCAATAGACAAGCATCATTCCAAGAACGATATCCCTTACCAT
27[126] CGCACTCATCGAGACAAGCAATGAAACCTCAGACTGCCCCCT
28[104] CGGGAGAAGCCTTTAGAGATCGACTTTTCTCATCTACTCAGG
28[118] CTGTAATGGTCATTACAGAGGTTATACCCCGGAATGGGATAG
29[105] GCTATCAACTTTTGACCAAAAACATTATACTAATATAGCAAC
29[140] ATCCTAAGTTTATCAACAATAAAGTAATGTAAATAGAAAATT
30[104] AACAAGAGAATCGAAAGATTGATTTGTAAATCATACAGGCGC
31[112] TAAATTGGCGAAACACCAACTTTGAACAACCGTGGGAACAAA
32[125] AGTACAACGTTAATCGTCGGATTCTCAGTAGGGCTAATATAA
32[139] ATAAGAGTAATTGAAAAGCCACCGGAATATAAGGCAATATAT
33[91] AAATGTGTAATGGGCCTCAGGGCAAAGCGGTCGACTTTCCTG
34[104] TTGACCGAGCGAGTAACAACCATTTTGTAAATATTTGGAGCA
34[146] CCAGTATGAATCGCCATATTTTTTTCGAAGACGACCGCGCCT
34[153] ATTCTTATTTACCATTGAGGGCACCGACCACCAGTGCAGCAC
35[112] GCCAGCTTTCCGGAATAAACAACGCTCAATCAATATTGACGG
36[76] AGGCTGCTGCCAAGCTTGCATGCCTGCAGCCATTCGAAACAC
36[118] GCTTCTGGGGTACCGAGCTACTTTTTCAGTTAAATAGTATGT
37[105] GATCCCCGTGCCGGAATTGGGAACGGTGATTTTCAAGGTGTA
37[140] TTTAGTTAATTTCACGACCGTAATACATAAGACACACACTGA
38[139] CAAAGAACGCGAGATCCGGCTCCTTATTTACCAGACGCCTGT
38[154] AATCCAATTAACTATAGAACTGACGCAATCGTCACCTCAGCGG
39[119] ATTAACCAACGAATTCGTAATCATGGTCGTGAGCTTAATCAT
41[112] GCTGATTGCCCTGAAATTGCGTTGCGCTCACTGCCGGCAACA
41[126] TAGCTTACCGAACAAGCAAGAGAATTGAATTAACTACAGGGA
41[133] GATTAAGTCCTTGAAAACATATATGTGAAATGGAATGATGAA
42[104] GAGTTGCTGAGACGCGCTTTCCTAATGAATAGCTGTCTAGAG
42[111] CCTGAGAAAGAATAGCCCGAAATCAATAGCTCACCGCCTGGC
42[156] TTAATTTTCCTTCTGTATTCATTTAACAAAACGCGCAGTGCTTTG
43[140] ACCTTGCCTTAGAAACGCTGAGGTCTGAGGTTATACGCAAGA
44[90] TGGAACACATCACTTGCCTGAGTAGAAGGTGTTGTCAAAATATTAGGAA
45[112] ACTATAAAAGAAGAACAGTACTAATAAGAAAAGATTCATCAG
46[132] TTTCAATACAATAAAACAGTACCTTTTAATATCAAAGCGCAT
46[139] TTATTCAACAAACATCAAGAAGAATTACAAATAGCAAGTAAG
47[112] TTGGCTCGGGAGAATACCTGACGGGAGAGTTAAGCGGATTTTAGCATTC
48[104] CACACGATATTAGTCTTTACCGAGAGTAATCTCCACGACAAT
48[153] AACGTCAGATGAATAAAACAGAGAATAAATTTTTTACAACTA
49[91] GAATGGCCCAGTAAGTCTGAAATGGATTTAATATCAACGAGA
49[126] AATTATTGAAGGGTTAGAACCTATCATCCATATTAGCACCCA
49[133] TGCACGTATACAGTCGGATTCGCCTGATAGGCGAACTGAACA
51[105] GTGCCACGCTGAATTAAACTGATAGCCCTAAAACATGCAACA
51[140] CAGATGAGCCAGTTCTGAATCTTTAGCGATCGATAAGCACCA
51[147] TGGCAATCATTTTGCGGAACACCGAACGTGAAGCCAGAACGC
52[90] AGCATCACCTTGCTATTTGAG
52[104] AATGAAAAGACTTTAGTTGATGAGCTGACATTAACCGGTTGT

| | |
|---|---|
| 52[125] | AGGAGCGAACTCGTGTTGCTATTAGGCTTATCCGGTATTCTA |
| 18[63] | ATACATAACGCCAAATCATAAGGATAGC |
| 21[62] | TTGAATCATTGCATCAAAAAGAAGCGAA |
| 17[76] | CCCCTCAAATGCTTAAAATGTTTAGACTCCCTCGT |
| 49[83] | ATATTACCTCAATCTAAAAGGGACATTCCAGACAA |

**Table 2. Table of unfunctionalized antibody capture staples**

| Name | Sequence |
|---|---|
| 29[147] | TTTACGATTCCTTAAGCCGTTTTTATTTTCATC |
| 32[169] | AATTTAGGCAGAGGCAAACAACGCCAACATGT |
| 44[162] | TTTAACAATAATCGTCGC |
| 50[165] | TCCTGATTGTTTGAAATTGCGTAG |

**Table 3. Table of functionalized antibody capture staples**

| Name | Sequence |
|---|---|
| 29[147] | TTTACGATTCCTTAAGCCGTTTTTATTTTCATCAGTCGAAGAGCACTAGGTAGAG |
| 32[169] | AATTTAGGCAGAGGCAAACAACGCCAACATGTAGTCGAAGAGCACTAGGTAGAG |
| 44[162] | TTTAACAATAATCGTCGC AGTCGAAGAGCACTAGGTAGAG |
| 50[165] | TCCTGATTGTTTGAAATTGCGTAGAGTCGAAGAGCACTAGGTAGAG |

**Table 4. Table of unfunctionalized cholesterol capture staples**

| Name | Sequence |
|---|---|
| 8[169] | TTAGAGCCAGCAAAATTTGAGCCATTTGGGAA |
| 10[169] | GGCGACATTCAACCGAGCGCCAAAGACAAAAG |
| 12[169] | ATATAAAAGAAACGCAACATAAAGGTGGCAAC |
| 14[162] | ATACCCAAAATGTAAATG |
| 16[162] | AAGCCCTTTCAATAGTGA |
| 18[162] | GAGCGCTAAATCTTACCG |
| 20[165] | AATAGCAGCCTTAGGGTAATT |
| 22[165] | TCTTTCCAGAGCGTCAAAAATGAA |
| 24[165] | CCCGACTTGCGGCGCTAACGAGCG |
| 26[153] | ACCGCGCGAGGCGTTTTAGCGAACCT |
| 26[165] | GTAGGAATCATTCGTAATCAGTAG |
| 28[169] | CAATAATCGGCTGTCTGCATGTAGAAACCAAT |
| 30[169] | TTCAGCTAATGCAGAAGACAATAAACAACATG |
| 34[169] | CATATGCGTTATACAAAAAGCCTGTTTAGTAT |
| 36[169] | TAATGGTTTGAAATACTCTTCTGACCTAAATT |
| 40[162] | ATTTATCAAAATCATAGAAGAGTTTAAGAAAATAGCT |
| 48[165] | ATTTTCAGGTTTAATACCAAG |
| 52[165] | GAGTAACATTATTCATCAATATAA |

**Table 5. Table of functionalized cholesterol capture staples**

| Name | Sequence |
|---|---|
| 8[169] | TTTTT TTAGAGCCAGCAAAATTTGAGCCATTTGGGAA TGACAGGATTAGCAGAGCGAGG |
| 10[169] | TTTTT GGCGACATTCAACCGAGCGCCAAAGACAAAAG TGACAGGATTAGCAGAGCGAGG |
| 12[169] | TTTTT ATATAAAAGAAACGCAACATAAAGGTGGCAAC TGACAGGATTAGCAGAGCGAGG |
| 14[162] | TTTTT ATACCCAAAATGTAAATG TGACAGGATTAGCAGAGCGAGG |
| 16[162] | TTTTT AAGCCCTTTCAATAGTGA TGACAGGATTAGCAGAGCGAGG |
| 18[162] | TTTTT GAGCGCTAAATCTTACCG TGACAGGATTAGCAGAGCGAGG |
| 20[165] | TTTTT AATAGCAGCCTTAGGGTAATT TGACAGGATTAGCAGAGCGAGG |
| 22[165] | TTTTT TCTTTCCAGAGCGTCAAAAATGAA TGACAGGATTAGCAGAGCGAGG |
| 24[165] | TTTTT CCCGACTTGCGGCGCTAACGAGCG TGACAGGATTAGCAGAGCGAGG |
| 26[153] | TTTTT ACCGCGCGAGGCGTTTTAGCGAACCT TGACAGGATTAGCAGAGCGAGG |
| 26[165] | TTTTT GTAGGAATCATTCGTAATCAGTAG TGACAGGATTAGCAGAGCGAGG |
| 28[169] | TTTTT CAATAATCGGCTGTCTGCATGTAGAAACCAAT TGACAGGATTAGCAGAGCGAGG |
| 30[169] | TTTTT TTCAGCTAATGCAGAAGACAATAAACAACATG TGACAGGATTAGCAGAGCGAGG |
| 34[169] | TTTTT CATATGCGTTATACAAAAAGCCTGTTTAGTAT TGACAGGATTAGCAGAGCGAGG |
| 36[169] | TTTTT TAATGGTTTGAAATACTCTTCTGACCTAAATT TGACAGGATTAGCAGAGCGAGG |
| 40[162] | TTTTT ATTTATCAAAATCATAGAAGAGTTTAAGAAAATAGCT TGACAGGATTAGCAGAGCGAGG |
| 48[165] | TTTTT ATTTTCAGGTTTAATACCAAG TGACAGGATTAGCAGAGCGAGG |
| 52[165] | TTTTT GAGTAACATTATTCATCAATATAA TGACAGGATTAGCAGAGCGAGG |

**Table 6. Table of unfunctionalized reporter-dye capture staples at the needle base**

| Name | Sequence |
|---|---|
| 25[84] | TTCATTTGGGGCGCTCCCAATACTAAAGTTTTGCG |
| 29[84] | ATTTTTGATTTCAACGCAAGGATAAAAACGGAGAG |
| 37[56] | AAAACGACGGCCAGGCAACTGGAATAAGAAGAGTA |
| 45[70] | TTGATTAGTAATAAAGAGTCCTTACCAGAATGCAG |
| 53[84] | GATTTAGAAGTATTAATCTAACACCGCCTCGCCAT |

**Table 7. Table of functionalized reporter-dye capture staples at the needle base**

| Name | Sequence |
|---|---|
| 25[84] | TTCATTTGGGGCGCTCCCAATACTAAAGTTTTGCG TGGCTGTGGTGTGCGAGTAACT |
| 29[84] | ATTTTTGATTTCAACGCAAGGATAAAAACGGAGAG TGGCTGTGGTGTGCGAGTAACT |
| 53[84] | AAAACGACGGCCAGGCAACTGGAATAAGAAGAGTA TGGCTGTGGTGTGCGAGTAACT |
| 45[70] | TTGATTAGTAATAAAGAGTCCTTACCAGAATGCAG TGGCTGTGGTGTGCGAGTAACT |
| 37[56] | GATTTAGAAGTATTAATCTAACACCGCCTCGCCAT TGGCTGTGGTGTGCGAGTAACT |

**Table 8. Table of unfunctionalized payload-dye strands at the needle apex**

| Name | Sequence |
|---|---|

| | |
|---|---|
| 0[195] | AGAAGGAACCACCGGAACCGCAGAGCCGTTCACAA |
| 3[203] | ACAAATAGTAAGCGCCCCCTGCCTATTTAAGAGGCAGAGCCG |
| 4[195] | CCACCAGAACCACCCAGAGCCTTAGGAT |
| 4[216] | GAGCCACCACCCTCCTCCCTCTGAGACT |

**Table 9. Table of functionalized payload-dye strands at the needle apex**

Here Cy3 or SeTau647N was used as the fluorophore.

| Name | Sequence |
|---|---|
| 0[195] | Fluorophore-S-S-AGAAGGAACCACCGGAACCGCAGAGCCGTTCACAA |
| 3[203] | Fluorophore-S-S-ACAAATAGTAAGCGCCCCCTGCCTATTTAAGAGGCAGAGCCG |
| 4[195] | Fluorophore-S-S-CCACCAGAACCACCCAGAGCCTTAGGAT |
| 4[216] | Fluorophore-S-S-GAGCCACCACCCTCCTCCCTCTGAGACT |

**Table 10. Table of other sequences**

Here, Cy5 and SeTau647N was used as the fluorophore.

| Name | Sequence |
|---|---|
| Antibody strand | /C6NH2/CTC TAC CTA GTG CTC TTC GACT |
| Cholesterol strand | CCT CGC TCT GCT AAT CCT GTC A - Cholesterol |
| Reporter-dye fluorophore | /5AmMC6/AGTTACTCGCACACCACAGCCA |
